# A low sugar diet enhances *Drosophila* body size in males and females via sex-specific mechanisms

**DOI:** 10.1101/2021.03.03.433819

**Authors:** Jason W. Millington, Lianna W. Wat, Ziwei Sun, Paige J. Basner-Collins, George P. Brownrigg, Elizabeth J. Rideout

## Abstract

In *Drosophila*, changes to dietary protein elicit different body size responses between the sexes. Whether this sex difference in nutrient-dependent body size regulation extends to other nutrients, such as dietary sugar, remains unclear. Here, we show that reducing dietary sugar enhanced body size in *Drosophila* male and female larvae. Indeed, the largest body size was found in larvae reared in a diet without added sugar. Despite the equivalent body size effects of a low sugar diet between males and females, we detected sex-specific changes to the insulin/insulin-like growth factor (IIS) and target of rapamycin (TOR) signaling pathways. Further, we show that the metabolic changes observed in larvae reared on a low sugar diet differ between the sexes. Thus, despite identical phenotypic responses to dietary sugar in males and females, distinct changes to cell signaling pathways and whole-body metabolism were associated with the increased body size in each sex. This highlights the importance of including both sexes in all mechanistic studies on larval growth, as males and females may use different molecular and metabolic mechanisms to achieve similar phenotypic outcomes.

## INTRODUCTION

In *Drosophila*, dietary nutrients impact the rate and duration of larval growth to influence final body size. Nutrient quantity promotes growth during larval development, as conditions where nutrients are plentiful favour larger body sizes (Edgar, 2006; Hietakangas and Cohen, 2009; Nijhout et al., 2014). Nutrient quality is also critical in regulating larval growth, as individual macronutrients differ in their body size effects. For example, while dietary protein promotes a larger body size across a wide concentration range (Britton and Edgar, 1998; Britton et al., 2002; Edgar, 2006; Shingleton et al., 2017), moderate or high levels of dietary sugar inhibit growth and reduce body size (Musselman et al., 2011; Pasco and Léopold, 2012; Reis, 2016). This suggests a complex relationship between individual macronutrients and body size.

One factor that influences the magnitude of nutrient-dependent changes to *Drosophila* body size is biological sex (McDonald et al., 2020 preprint; Millington et al., 2021a; Shingleton et al., 2017; Stillwell et al., 2010; Teder and Tammaru, 2005). For example, manipulating nutrient quantity by altering dietary protein and carbohydrates causes sex-biased trait size effects (Shingleton et al., 2017). Male and female phenotypic responses to altered nutrient quality also differ, as the magnitude of protein-dependent changes to body size are larger in females (Millington et al., 2021a). Due to the widespread use of mixed-sex groups in larval growth studies, however, it remains unclear whether sex-specific body size responses to dietary protein extend to other macronutrients, such as sugar.

Our examination of larval development revealed that a reduction in dietary sugar significantly increased the rate of growth and body size in males and females. Indeed, the largest body size was observed in a diet with no added sugar. Despite the equivalent body size increase in males and females, sex-specific mechanisms underlie the larger body size of larvae raised in a low sugar diet. In females, the low sugar diet stimulated increased target of rapamycin (TOR) pathway activity, whereas the activity of the insulin/insulin-like growth factor signaling pathway (IIS) was enhanced in males. Genetic studies confirmed that these female- and male-specific changes to TOR and IIS, respectively, were important for the low sugar-induced increase in body size, and biochemical studies revealed sex-specific changes to metabolic gene expression and metabolism. Together, our findings provide additional mechanistic insight into how dietary sugar affects development by revealing sex-specific changes to cell signaling pathways and metabolism. This highlights the importance of including both sexes in all larval growth studies, as we show that equivalent phenotypic outcomes may be achieved via distinct mechanisms in each sex.

## RESULTS AND DISCUSSION

### A low sugar diet promotes an increased rate of growth and augments body size

To determine the body size effects of dietary sugar in each sex, we quantified pupal volume in *white^1118^* (*w*; FBgn0003996) male and female larvae reared in diets with different levels of dietary sugar. Because dietary sugar represses growth in a mixed-sex larval group (Musselman et al., 2011; Pasco and Léopold, 2012), we started with a widely-used diet (1S) (Lewis, 1960) and removed sugar in a stepwise manner until no added sugar remained (0S). In *w^1118^* females, body size was significantly larger in larvae cultured on a diet with half (0.5S), or one-quarter (0.25S), the amount of sugar found in 1S (Fig. 1A). Interestingly, the largest body size was found in female larvae reared in 0S (Fig. 1A).

**Figure 1.**
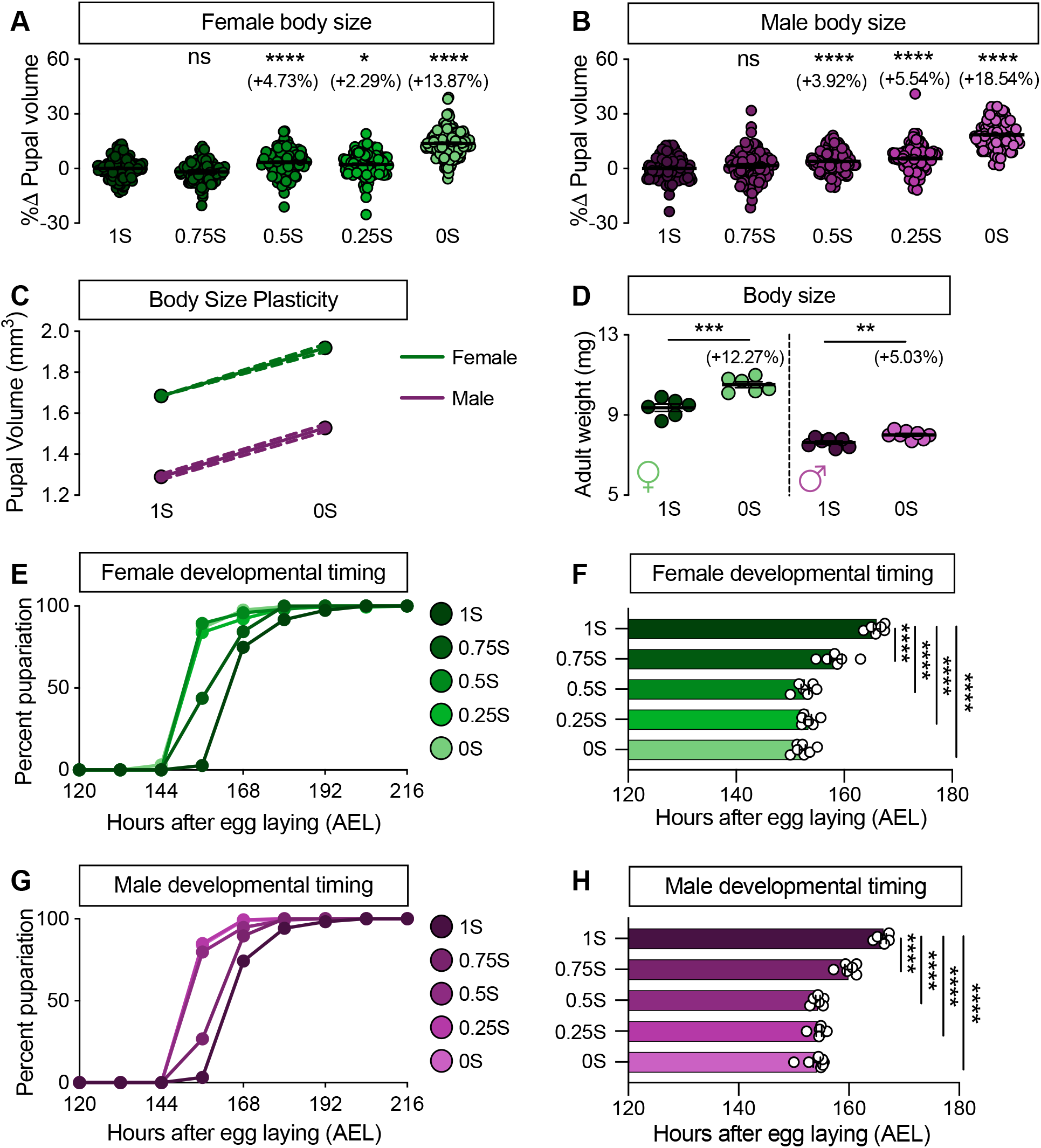
A low sugar diet promotes an increased rate of growth and final body size. (A) Pupal volume in *w^1118^* females cultured on 1S, 0.75S, 0.5S, 0.25S, and 0S. (B) Pupal volume in *w^1118^* males cultured on 1S, 0.75S, 0.5S, 0.25S, and 0S. (C) Reaction norms for pupal volume in both sexes plotted using 1S and 0S data from A and B. (D) Adult weight in *w^1118^* female and male flies reared on 1S and 0S. (E, F) Time to pupariation in *w^1118^* females cultured on 1S, 0.75S, 0.5S, 0.25S, and 0S. (G, H) Time to pupariation in *w^1118^* males cultured on 1S, 0.75S, 0.5S, 0.25S, and 0S. * p<0.05; ** p<0.01; *** p<0.001; **** p<0.0001; ns indicates not significant; error bars indicate SEM; dashed lines indicate 95% confidence interval. *p*-values, samples sizes, and statistical tests are in Supplementary file 4.

In *w^1118^* males, body size was significantly larger in larvae reared on 0.5S and 0.25S compared with larvae raised on 1S (Fig. 1B). As in females, the largest body size among males was recorded on 0S (Fig. 1B). Importantly, the body size effects of reduced sugar diets were equivalent between the sexes (Fig. 1C; Supplemental file 4), a finding we reproduced using adult weight (Fig. 1D), indicating that phenotypic responses to dietary sugar were not different between males and females. Because a diet with fewer calories has no effect on body size (Millington et al., 2021a), our findings suggest that the larger size of larvae raised in 0S can be attributed to less dietary sugar. This agrees with data from a mixed-sex larval group showing that dietary sugar inhibits growth (Musselman et al., 2011; Pasco and Léopold, 2012), and extends previous findings by showing the body size effects occur in both sexes.

To determine the growth rate in larvae reared on low sugar diets, we measured the time between egg-laying and pupariation in both sexes. In *w^1118^* females, time to 50% pupariation was significantly shorter in larvae raised on each reduced-sugar diet compared with genotype-matched larvae cultured in 1S (Fig. 1E, F). Time to 50% pupariation was also reduced in *w^1118^* males raised on each reduced-sugar diet compared with genotype-matched males reared on 1S (Fig. 1G, H). Given that diets with less added sugar shorten the larval growth period and increase body size, our data suggests the larval growth rate in each sex was significantly accelerated in a low sugar context.

### A low sugar diet has sex-biased effects on insulin/insulin-like growth factor (IIS) and target of rapamycin (TOR) signaling

Many signaling pathways control organismal and tissue growth during development, however, IIS and TOR have emerged as key regulators of nutrient-dependent growth (Gokhale and Shingleton, 2015; Grewal, 2009; Koyama and Mirth, 2018; Lecuit and Le Goff, 2007; Teleman, 2010). Indeed, high levels of IIS and TOR activity promote a larger body size (Böhni et al., 1999; Britton et al., 2002; Chen et al., 1996; Fernandez et al., 1995; Patel et al., 2003; Poltilove et al., 2000; Zhang et al., 2000). Given the larger body size of males and females cultured on 0S, we examined IIS and TOR activity in larvae reared on 0S and 1S. To measure IIS activity, we quantified mRNA levels of genes that are coregulated by transcription factor Forkhead box, sub-group O (Foxo; FBgn0038197) (*e.g. Insulin receptor* (*InR*; FBgn0283499), *brummer* (*bmm;* FBgn0036449), and *eukaryotic initiation factor 4E-binding protein* (*4E-BP*; FBgn0261560)). For example, when IIS activity is high, Foxo is repressed and mRNA levels of *InR*, *bmm*, and *4E-BP* are low (Alic et al., 2011; Jünger et al., 2003; Puig and Tjian, 2005; Zinke et al., 2002).

In *w^1118^* females, mRNA levels of Foxo target genes were not different between larvae reared in 0S and 1S (Fig. 2A), suggesting IIS activity was not altered in females. In contrast, mRNA levels of Foxo target genes were significantly lower in *w^1118^* male larvae in 0S (Fig. 2B), indicating enhanced IIS activity. Importantly, feeding behaviour was not different between the sexes in either diet (Fig. 2C). While increased IIS activity in males raised on 0S may be due to improved insulin sensitivity, changes to mRNA levels of two genes upregulated by insulin insensitivity (*Neural Lazarillo* (*NLaz*; FBgn0053126) and *puckered* (*puc*; FBgn0243512)) were not consistent with improved insulin sensitivity in either sex (Fig. 2D, E) (Lourido et al., 2021; Pasco and Léopold, 2012). Indeed, altered *puc* mRNA levels in males likely reflect Foxo activity, as *puc* is a Foxo target (Bai et al., 2013). Together, these findings reveal a previously unrecognized sex difference in IIS regulation in a low sugar context, adding to a growing body of evidence showing sex differences in the nutrient-dependent regulation of IIS in larvae (Millington et al., 2021a).

**Figure 2.**
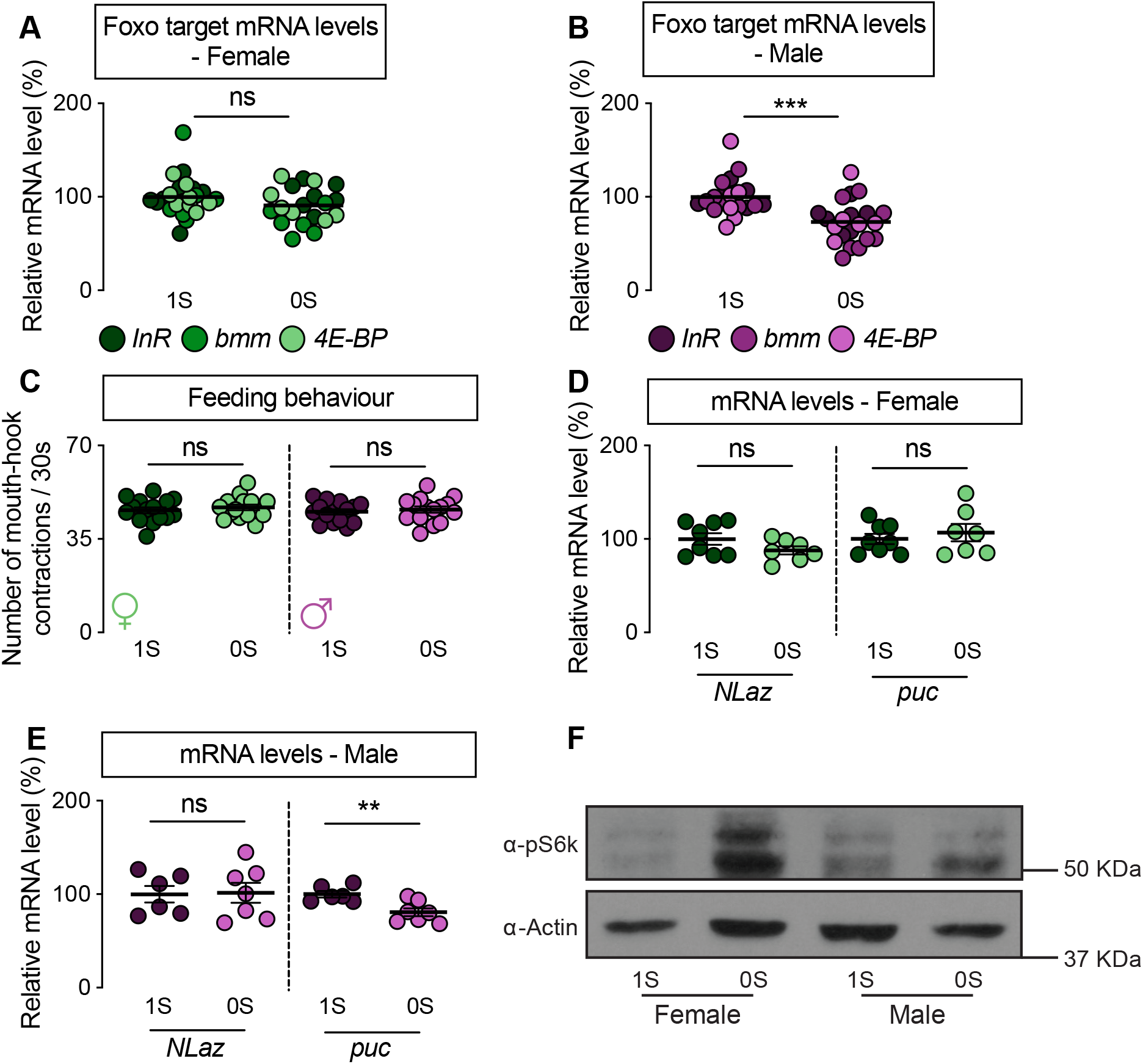
A low sugar diet has sex-biased effects on insulin/insulin-like growth factor (IIS) and target of rapamycin (TOR) signaling. (A) mRNA levels of Foxo target genes (*InR*, *bmm*, and *4E-BP*) in female larvae reared on 1S or 0S. (B) mRNA levels of Foxo target genes in male larvae reared on 1S or 0S. (C) Mouth-hook contractions in *w^1118^* female and male larvae raised on 1S or 0S. (D) mRNA levels of *NLaz* and *puc* in female larvae raised in 1S or 0S. (E) mRNA levels of *NLaz* and *puc* in male larvae raised in 1S or 0S. (F) Levels of phosphorylated S6 kinase (pS6k) in females and males raised in 1S or 0S. ** p<0.01; *** p<0.001; ns indicates not significant; error bars indicate SEM. *p*-values, samples sizes, and statistical tests are in Supplementary file 4.

We next measured TOR activity by monitoring the phosphorylation of TOR’s downstream target Ribosomal protein S6 kinase (S6k; FBgn0283472). In *w^1118^* females, levels of phosphorylated S6k (pS6k) were higher in 0S in multiple biological replicates (Fig. 2F; Fig S1A), an effect we did not reproduce in *w^1118^* males (Fig. 2F; Fig. S1A). This suggests that the low sugar diet caused a female-biased increase in TOR activity, revealing a previously unrecognized sex difference in the nutrient-dependent regulation of TOR. Taken together, these findings not only extend knowledge of sex-specific IIS regulation, but also provide the first report of sex-biased TOR regulation.

### Sex-biased requirement for IIS, *Drosophila* insulin-like peptides, and TOR in promoting the low sugar-induced increase in body size

To determine whether sex-biased changes to IIS and TOR play a role in mediating the low sugar-induced increase in body size, we measured body size in male and female larvae carrying mutations in each pathway that blunt high levels of IIS and TOR activation (Chen et al., 1996; Millington et al., 2021a; Rideout et al., 2015; Zhang et al., 2000). To determine the requirement for IIS, we measured pupal volume in *w^1118^* larvae, and in larvae heterozygous for a hypomorphic allele of *InR* (*InR^E19^/+*), in 1S and 0S. While body size was larger in *w^1118^* male larvae reared on 0S than 1S (Fig. 3A), the low sugar-induced increase in body size was blocked in *InR^E19^/+* males (Fig. 3A; genotype:diet interaction *p*<0.0001). Thus, IIS activity was required in males for increased body size in 0S. In contrast, female *w^1118^* and *InR^E19^/+* larvae reared on 0S were significantly larger than genotype-matched females raised on 1S (Fig. 3B). While we detected a significant genotype:diet interaction in females (*p*<0.0001), the magnitude of genotype effects on the body size response to low sugar was smaller in females than in males (sex:diet:genotype interaction *p*=0.0114). Thus, reduced IIS function had a male-biased impact on the low sugar-induced increase in body size.

**Figure 3.**
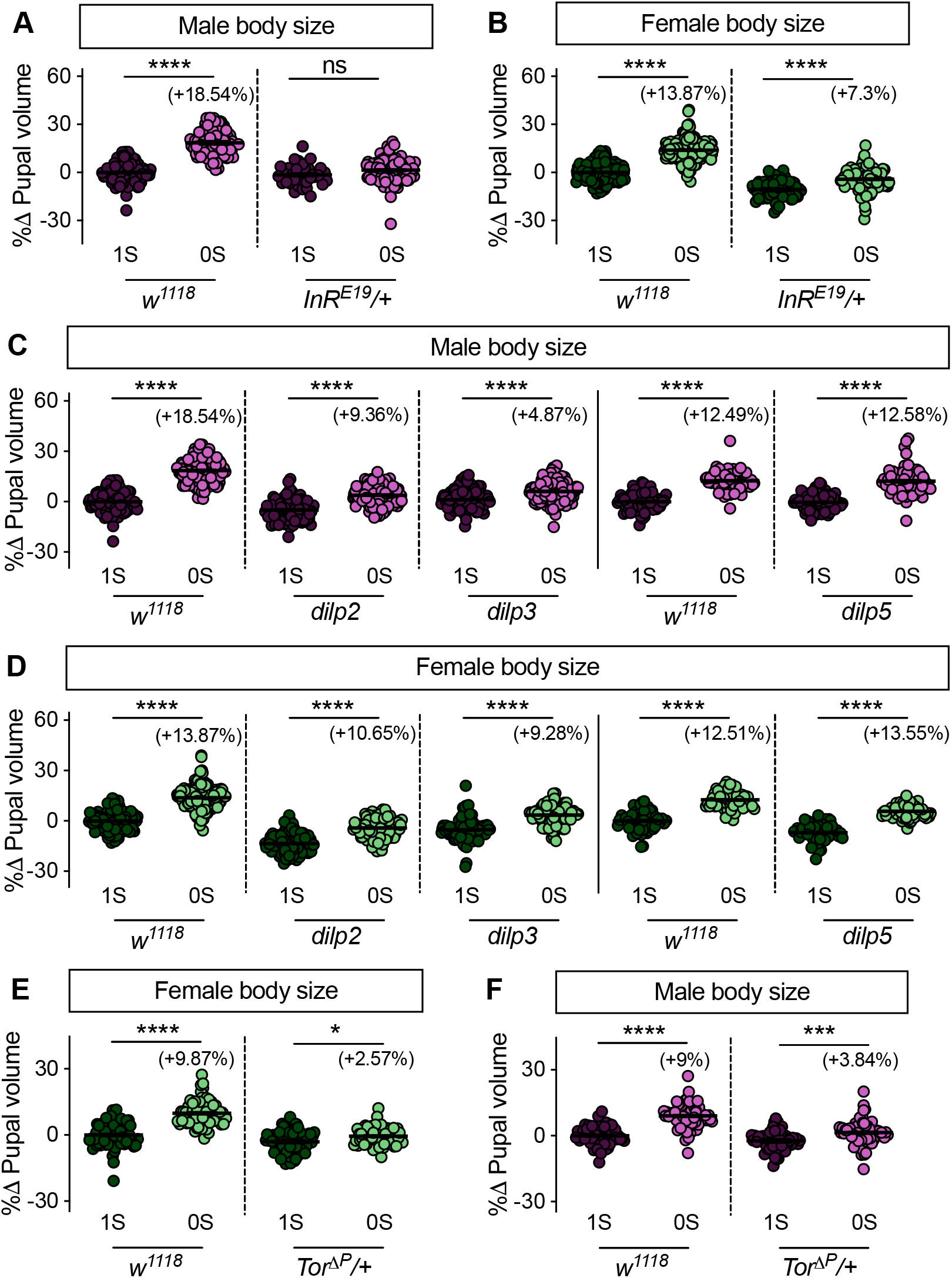
Sex-biased requirement for IIS, *Drosophila* insulin-like peptides, and target of rapamycin (TOR) in promoting the low sugar-induced increase in body size. (A) Pupal volume in *w^1118^* and *InR^E19^/+* males cultured on 1S or 0S. (B) Pupal volume in *w^1118^* and *InR^E19^/+* females cultured on 1S or 0S. (C) Pupal volume in *w^1118^*, *dilp2* mutant, *dilp3* mutant, and *dilp5* mutant males reared on 1S or 0S. (D) Pupal volume in *w^1118^*, *dilp2* mutant, *dilp3* mutant, and *dilp5* mutant females reared on 1S or 0S. (E) Pupal volume in *w^1118^* and *Tor^ΔP^/+* females cultured on 1S or 0S. (F) Pupal volume in *w^1118^* and *Tor^ΔP^/+* males cultured on 1S or 0S. * p<0.05; *** p<0.001; **** p<0.0001; ns indicates not significant; error bars indicate SEM. *p*-values, samples sizes, and statistical tests are in Supplementary file 4. Note: parallel collection of multiple genotypes and diets means that *w^1118^* control data in 0S and 1S are the same in Fig. 1A, B, 3A-D.

Beyond InR, we reared male and female larvae lacking the coding sequences for *Drosophila insulin-like peptide 2* (*dilp2*; Fbgn0036046), *Drosophila insulin-like peptide 3* (*dilp3*; Fbgn0044050), and *Drosophila insulin-like peptide 5* (*dilp5*; Fbgn0044038) on 0S and 1S (Grönke et al., 2010). These Dilps are produced and secreted by insulin-producing cells in the brain (Brogiolo et al., 2001; Géminard et al., 2009; Ikeya et al., 2002; Rulifson et al., 2002). Circulating Dilps stimulate IIS activity and growth by binding to InR on target cells (Teleman, 2010). In males, loss of *dilp2* and *dilp3* blunted the low sugar-induced increase in body compared with *w^1118^* controls (genotype:diet *p*<0.0001 for both); loss of *dilp5* had no effect (genotype:diet *p*=0.9751) (Fig. 3C). In females, while loss of *dilp2* and *dilp3* (genotype:diet *p*<0.0001 for both), but not *dilp5* (genotype:diet *p*=0.9389), blunted the low sugar-induced increase in body size (Fig. 3D), the magnitude of genotype effects on the increase in body size were larger in males for *dilp3* (sex:diet:genotype interaction: *p=*0.0003), with a similar trend in *dilp2* (sex:diet:genotype interaction: *p=*0.0627). Thus, we identify a male-biased requirement for several genes that influence IIS activity in regulating the low sugar-induced increase in body size, a finding that aligns with the male-specific increase in IIS activity in 0S.

To determine the requirement for TOR in mediating the low sugar-induced increase in body size, we measured pupal volume in larvae heterozygous for a hypomorphic allele of *Target of rapamycin* (*Tor*; *Tor*^Δ*P*^/+) (Zhang et al., 2000). In *w^1118^* females, larvae reared on 0S were significantly larger than genotype-matched larvae raised on 1S (Fig. 3E); however, this low sugar-induced increase in body size was blunted in *Tor*^Δ*P*^/+ female larvae (Fig. 3E; genotype:diet interaction *p*<0.0001). This suggests the low sugar-induced increase in TOR activity in females was required to achieve a larger body size. In *w^1118^* and *Tor*^Δ*P*^/+ males, pupal volume was significantly larger in larvae raised on 0S compared with genotype-matched larvae reared on 1S (Fig. 3F). While the low sugar-induced increase in body size was smaller in *Tor*^Δ*P*^/+ males compared with controls (genotype:diet *p*<0.0001), the magnitude of genotype effects on the body size response were larger in females than in males (sex:diet:genotype interaction (*p*=0.0303). This reveals a previously unrecognized female-biased requirement for TOR activity in regulating body size in a low sugar context.

### A low sugar diet has sex-specific effects on metabolic gene expression and whole-body metabolism

IIS and TOR promote increased body size by regulating diverse aspects of metabolism, (*e.g*. triglyceride storage, protein synthesis, glucose homeostasis) (Grewal, 2009; Musselman and Kühnlein, 2018; Teleman et al., 2008). We therefore measured mRNA levels of a selection of genes implicated in metabolic regulation. In *w^1118^* male and female larvae reared on 0S, we found significant changes to mRNA levels compared with larvae reared on 1S, many of which were sex-specific (Fig. 4A). For genes encoding proteins involved in fat metabolism, 9/14 and 5/14 genes showed low sugar-induced changes to mRNA levels in males and females, respectively (Fig. 4A). For genes encoding ribosomal proteins, which play integral roles in protein synthesis, 1/12 and 6/12 genes showed low sugar-induced changes to mRNA levels in males and females, respectively (Fig. 4A). While this examination of mRNA levels includes only a fraction of genes that affect metabolism, this data suggests that a low sugar diet causes sex-biased changes to mRNA levels of metabolic genes.

**Figure 4.**
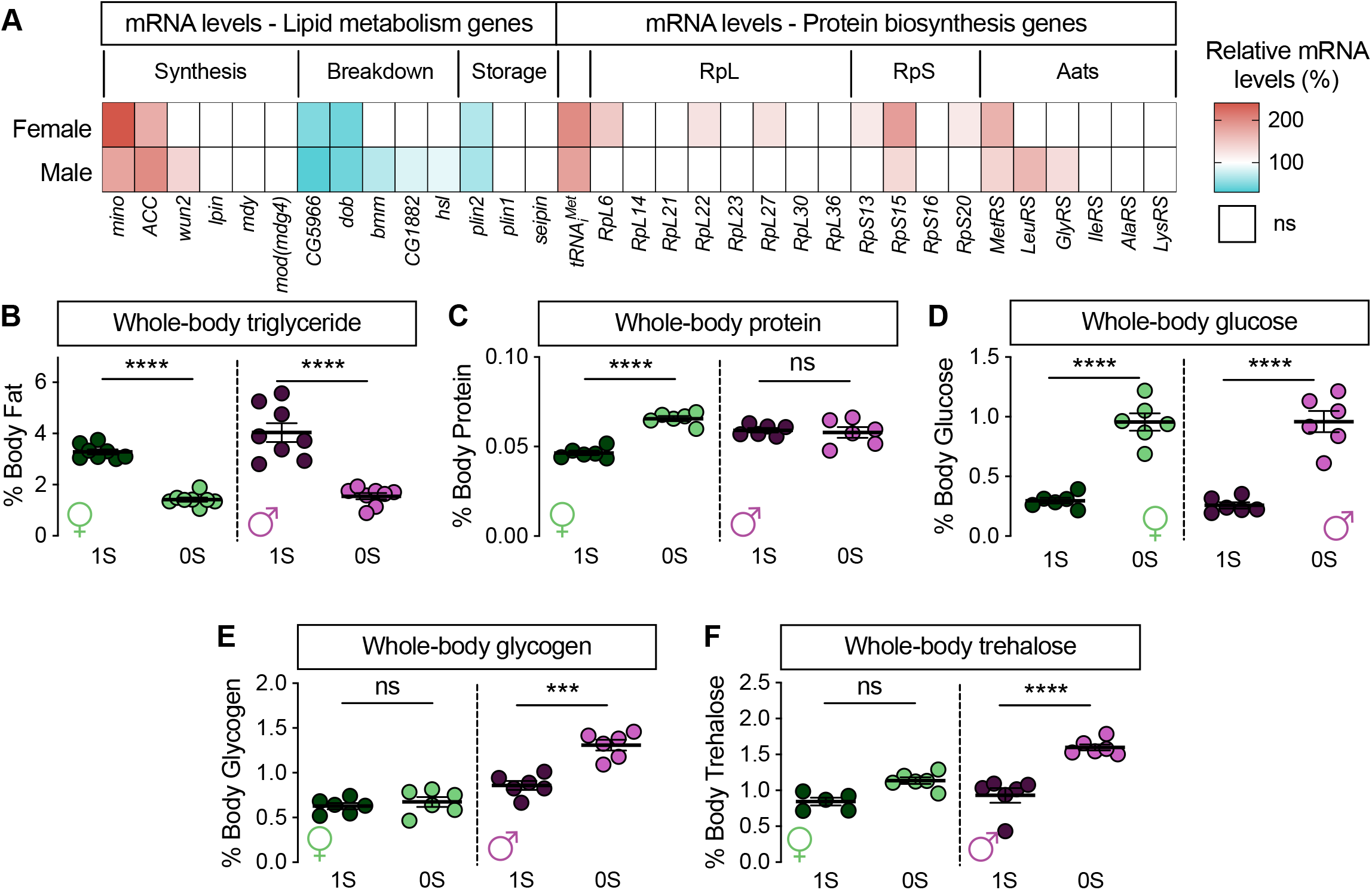
A low sugar diet has sex-biased effects on metabolic gene expression and metabolism. (A) mRNA levels of metabolic genes in *w^1118^* male and female larvae raised on 1S or 0S. (B) Whole-body triglyceride levels in *w^1118^* female and male larvae reared on 1S or 0S. (C) Whole-body protein levels in *w^1118^* female and male larvae raised on 1S or 0S. (D) Whole-body glucose levels in *w^1118^* female and male larvae cultured on 1S or 0S. (E) Whole-body glycogen levels in *w^1118^* female and male larvae reared on 1S or 0S. (F) Whole-body trehalose levels in *w^1118^* female and male larvae cultured on 1S or 0S. *** p<0.001; **** p<0.0001; ns indicates not significant; error bars indicate SEM. *p*-values, samples sizes, and statistical tests are in Supplementary file 4.

To determine the physiological significance of these sex-biased changes in mRNA levels (Gershman et al., 2007; Li et al., 2006; Mattila and Hietakangas, 2017; Teleman et al., 2008; Zinke et al., 2002), we measured whole-body levels of several macronutrients in male and female larvae reared in 0S and 1S. In both male and female *w^1118^* larvae reared on 0S, triglyceride levels were significantly reduced compared with sex-matched larvae cultured on 1S (Fig. 4B). Thus, a low sugar diet reduced adiposity in both sexes. In contrast, *w^1118^* females reared in 0S had significantly higher protein levels (Fig. 4C), an effect that was not reproduced in *w^1118^* males (Fig. 4C). This reveals a previously unrecognized sex difference in the regulation of whole-body protein in a low sugar context. While glucose levels were significantly higher in *w^1118^* male and female larvae raised in 0S (Fig. 4D), the low sugar diet caused a male-specific increase in glycogen and trehalose (Fig. 4E, F). Thus, we observed both sex-specific and non-sex-specific alterations in whole-body carbohydrate levels in a low sugar context, highlighting the importance of including both sexes when studying diet-induced metabolic changes. Indeed, our findings suggest sugar-induced changes to carbohydrate metabolism found in previous studies were possibly driven by effects in males (Musselman et al., 2011; Pasco and Léopold, 2012).

In conclusion, our study adds to a growing literature showing sex-specific effects of dietary nutrients on phenotypes such as body size, metabolism, lifespan, and fertility (Green and Extavour, 2014; Hudry et al., 2019; Klepsatel et al., 2020; Millington et al., 2021a; Regan et al., 2016; Wat et al., 2020). Because we show males and females activate distinct signaling pathways and exhibit sex-specific metabolic changes to achieve equivalent phenotypic outcomes, this suggests the absence of a sexually dimorphic phenotype in larval growth studies does not provide sufficient rationale for using single- or mixed-sex groups of animals. Instead, both sexes must be included to draw accurate conclusions regarding the signaling, metabolic, and body size effects of dietary nutrients.

## MATERIALS AND METHODS

### Data availability

Original images of pupae are available upon request. Raw values for all data collected and displayed in this manuscript are available in Supplementary File 1. All data necessary for confirming the conclusions of the article are present within the article, figures, tables, and supplementary files.

### Fly husbandry

Our 1S diet consists of 20.5 g/L sucrose, 70.9 g/L D-glucose, 48.5 g/L cornmeal, 45.3 g/L yeast, 4.55 g/L agar, 0.5g CaCl2•2H2O, 0.5 g MgSO4•7H2O, 11.77 mL acid mix (propionic acid/phosphoric acid). Our 0S diet consists of 48.5 g/L cornmeal, 45.3 g/L yeast, 4.55 g/L agar, 0.5g CaCl2•2H2O, 0.5 g MgSO4•7H2O, 11.77 mL acid mix. Details of 0.75S, 0.5S, and 0.25S diets can be found in Supplementary file 2. Larvae were raised at a density of 50 animals per 10 mL food at 25°C, and sexed by gonad size. Adult flies were maintained at a density of twenty flies per vial in single-sex groups.

### Fly strains

The following fly strains from the Bloomington *Drosophila* Stock Center were used: *w^1118^* (#3605), *InR^E19^* (#9646), *Tor*^Δ*P*^ (#7014). Additional fly strains include: *dilp2, dilp3,* and *dilp5* (Grönke et al., 2010). All fly strains were backcrossed for at least 6 generations, in addition to extensive prior backcrossing (Grönke et al., 2010; Millington et al., 2021a,b).

### Body size

Pupal volume and adult weight were measured as previously described (Delanoue et al., 2010; Millington et al., 2021a; Millington et al., 2021b; Rideout et al., 2015).

### Feeding behaviour

Feeding behavior was quantified as number of mouth-hook contractions per 30 s.

### Developmental timing

Time to pupariation was measured as previously described (Millington et al., 2021a). Time to 50% pupariation was calculated per replicate and used for quantification and statistical analysis.

### Metabolism assays

Each biological replicate consists of ten female or male larvae. Larvae were frozen on dry ice, and homogenized for lipid, protein, glucose, glycogen, and trehalose assays. All assays were performed as described in Tennessen et al. (2014) and Wat et al. (2020).

### RNA extraction and cDNA synthesis

RNA extraction and cDNA synthesis were performed as previously described (Marshall et al., 2012; Rideout et al., 2012; Rideout et al., 2015; Wat et al., 2020). Briefly, each biological replicate consists of ten *w^1118^* larvae frozen on dry ice and stored at −80°C. Each experiment contained 3-4 biological replicates per sex, and each experiment was performed at least twice. RNA was extracted using 500 μl Trizol (Thermo Fisher Scientific: #15596018) and precipitated using isopropanol and 75% ethanol. Pelleted RNA was resuspended in 200 μl molecular biology grade water (Corning, 46-000-CV) and stored at −80°C until use. For cDNA synthesis, an equal volume of RNA per reaction was DNase-treated and reverse transcribed using the QuantiTect Reverse Transcription Kit (Qiagen, 205314).

### Quantitative real-time PCR (qPCR)

qPCR was performed as previously described (Marshall et al., 2012; Rideout et al., 2012; Rideout et al., 2015; Wat et al., 2020). Primer list in Supplementary file 3.

### Preparation of protein samples, SDS-PAGE, and Western blotting

Samples were generated as previously described (Millington et al., 2021a). 20 μg of protein was loaded per lane, separated on a 12% SDS-PAGE gel in SDS running buffer, and transferred onto a nitrocellulose membrane (Bio-Rad) for 2 hr at 40 V on ice. Membranes were incubated for 24 hr in blocking buffer at 4°C (5% milk or 5% BSA in TBST 0.1%) and subsequently incubated with primary antibodies overnight at 4°C. Anti-pS6k (#9209, Cell Signaling), and anti-Actin (#8432, Santa Cruz) were used at 1:1000. After 3 x 2 min washes in 0.1% TBST, HRP-conjugated secondary antibodies were used at 1:5000 for pS6k (#65–6120; Invitrogen) and 1:3000 for actin (#7076; Cell Signaling). Membranes were washed (3 x 2 min, 2 x 15min) in 0.1% TBST, washed 1 x 5 min in TBS, and finally Pierce ECL was applied as per manufacturer’s instructions (#32134, Thermo Scientific)

### Statistical analysis

GraphPad Prism (GraphPad Prism version 8.4.3 for Mac OS X) was used for all statistical tests, and for figure preparation. Full details of statistical tests and *p*-values are listed in Supplementary file 4.

## ACKNOWLEDGEMENTS

We would like to thank Dr. Linda Partridge for sharing the *dilp2*, *dilp3*, and *dilp5* mutant strains. We thank the Bloomington *Drosophila* Stock Center (NIH P40OD018537) for stocks used in this study. We acknowledge critical resources and information provided by FlyBase (Thurmond et al., 2018); FlyBase is supported by a grant from the National Human Genome Research Institute at the U.S. National Institutes of Health (U41HG000739) and by the British Medical Research Council (MR/N030117/1). We acknowledge that our research takes place on the traditional, ancestral, and unceded territory of the Musqueam people; a privilege for which we are grateful.

---

**Figure S1.**
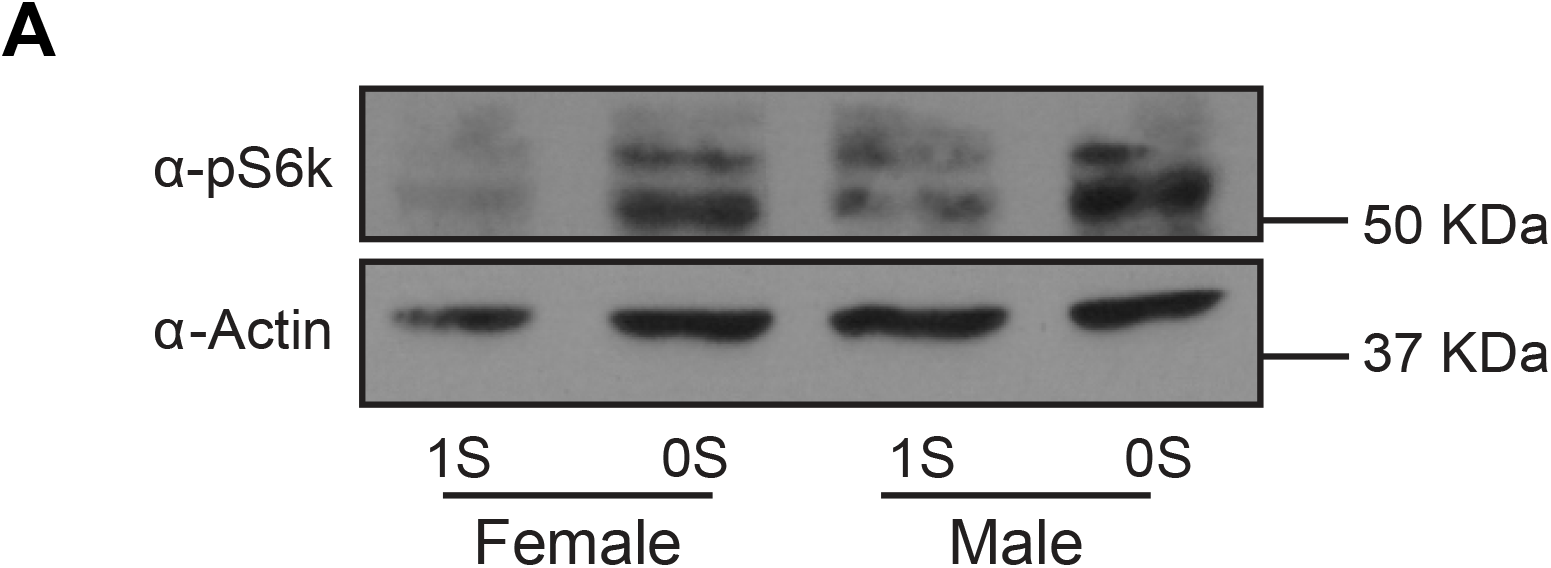
A low sugar diet has sex-biased effects on target of rapamycin (TOR) signaling. (A) Levels of pS6k in females and males raised in 1S or 0S.

